# Targeted metagenomic sequencing enhances the identification of pathogens associated with acute infection

**DOI:** 10.1101/716902

**Authors:** Cyndi Goh, Tanya Golubchik, M Azim Ansari, Mariateresa de Cesare, Amy Trebes, Ivo Elliott, David Bonsall, Paolo Piazza, Anthony Brown, Hubert Slawinski, Natalie Martin, Sylviane Defres, Mike J Griffiths, James E Bray, Martin C Maiden, Paula Hutton, Charles J Hinds, Tom Solomon, Ellie Barnes, Andrew J Pollard, Manish Sadarangani, Julian C Knight, Rory Bowden

## Abstract

The routine identification of pathogens during infection remains challenging because it relies on multiple modalities such as culture and nucleic acid amplification and tests that tend to be specific for very few of an enormous number of possible infectious agents. Metagenomics promises single-test identification, but shotgun sequencing remains unwieldy and expensive or in many cases insufficiently sensitive to detect the amount of pathogen material in a clinical sample. Here we present the validation and application of *Castanet*, a method for metagenomic sequencing with enrichment that exploits clinical knowledge to construct a broad panel of relevant organisms for detection at low cost with sensitivity comparable to PCR. *Castanet* targets both DNA and RNA, works with small sample volumes, and can be implemented in a high-throughput diagnostic setting. We used *Castanet* to analyse plasma samples from 573 patients from the GAinS sepsis cohort and CSF samples from 243 patients from the ChiMES meningitis cohort that had been evaluated using standard clinical microbiology methods, identifying relevant pathogens in many cases where no pathogen had previously been detected. *Castanet* is intended for use in defining the distribution of pathogens in samples, diseases and populations, for large-scale clinical studies and for verifying the performance of routine testing regimens. By providing sequence as output, *Castanet* combines pathogen identification directly with subtyping and phylo-epidemiology.

## Introduction

Many cases of acute infection, including meningitis and sepsis, are managed in the absence of a specific pathogen diagnosis in both high- and low-resource settings^1-5^. The primacy of empirical therapy reflects clinical urgency combined with the complexity of the testing landscape for acute conditions, a lack of assay sensitivity, and the time required to run a series of tests. A large number of potentially causative organisms must be considered, each with its own biology and consequently set of diagnostic tests. Even if all possible tests could be performed quickly enough to inform patient care, test sensitivity is limited by the low pathogen levels in many specimen types and by the progressive depletion of sample volume by each subsequent test. No single test is perfect: traditional culture methods are broadly applicable for bacteria but are slow and biased in favour of culturable and fast-growing organisms. Molecular tests such as organism-specific PCR are fast and specific, but risk false-negative results where variation among pathogen genomes leads to failure to detect the organism^6^.

Strategies to reduce the number of sequential tests and speed up pathogen detection have included the development of multi-pathogen assays such as multiplex PCRs^7-10^, panels of serological tests^11 12^, 16S DNA sequencing^13 14^, and most recently, metagenomic sequencing^15 16^. Diagnostic metagenomics – that is, deep sequencing of total nucleic acid directly from a clinical sample – is the only currently available technique that offers the promise of detecting the complete pathogen composition of a sample in a single assay, as well as information on virulence and antimicrobial susceptibility^15^. In practice however, diagnostic metagenomic applications are limited to sample types with high pathogen abundance^17 18^. In most readily available clinical samples, such as blood, the vast majority of sequences originate from the human host, since a single human cell contains as much nucleic acid as a thousand bacteria or up to a million viruses. In addition, material from commensal and contaminant organisms and kit reagents is essentially ubiquitous^19^, so a means is needed to distinguish pathogen sequences from background.

To be useful for routine clinical applications, metagenomics requires laboratory techniques that increase the relative yield of sequences that come from the pathogen(s) of interest. In efforts to harness the power of metagenomics for low-abundance samples, various purification or enrichment procedures have been employed to increase yield. For bacterial pathogens, the most common first step is culture, which can bias the result in favour of contaminating, culturable or fast-growing organisms. For viral pathogens and some bacteria, yield-maximising methods include low-speed centrifugation and/or filtration to remove host cells, sample treatment with nucleases to digest nucleic acid not protected within cells or virions^20 21^ and high-speed gradient centrifugation to concentrate virus particles (for review, see Duhaime and Sullivan 2012)^22^. Each of these procedures reduces throughput and may bring bias^23^.

An alternative and widely applicable approach is targeted enrichment of a subset of sequences in a metagenomic sample using nucleic acid probes, or “baits”. Targeted enrichment works at a range of scales from individual genes to the 50-60+ Mb of sequences defining the human exome^24-27^, and can be applied either to unprocessed clinical samples or to sequencing libraries. We previously demonstrated that enrichment is robust to substantial local divergence between probe and target sequences, making it possible to design multi-pathogen probe sets that simultaneously capture a diverse set of clinically relevant pathogen sequences^28^. Targeted enrichment for single or larger numbers of targeted organisms has been demonstrated in the study of viruses^28-32^ and bacteria^33 34^. However, significant practical hurdles remain in the application of sequence enrichment as a routine and comprehensive multi-pathogen test, including the development of efficient workflows, the need to balance the opposing considerations of the breadth of species detected and the cost and complexity of large probe panels^32 34 35^ and the requirement for bespoke analytical pathways.

Here we describe *Castanet*, a probe-based targeted enrichment methodology that can detect an arbitrarily wide selection of clinically important bacterial and viral pathogens, and interrogates both DNA and RNA sequences in the same workflow. *Castanet* is based on a clinically informed probe panel, designed to replace and augment first-line diagnostic tests for a range of acute infections, and is suitable for use in high-throughput settings. We validate and benchmark method performance both on a standard reference containing a pre-quantified mixture of known viruses^36^, and on clinical samples with well-quantified pathogen load. We then demonstrate the application of our method in clinical settings by using it to detect pathogens in patient samples from sepsis and meningitis, two clinically important, life-threatening syndromes where the available samples are notoriously low in pathogen load, and where conventional diagnostic testing often fails to find an infectious cause^15 37^.

## Results

### Design of probes for an extensive panel of disease-relevant pathogens

We compiled a list of viral and bacterial pathogens relevant to paediatric meningitis and adult sepsis from community-acquired pneumonia (CAP) in the UK (**Table 1**), adding several pathogens of current interest that were less likely to be present in UK samples. Considering the number of distinct entries on our list (116, from 17 virus families and 35 bacterial species), we inferred that any other organisms, including relevant fungal or parasite pathogens, would necessarily comprise rare causes of meningitis or pneumonia and sepsis.

**Table 1.**
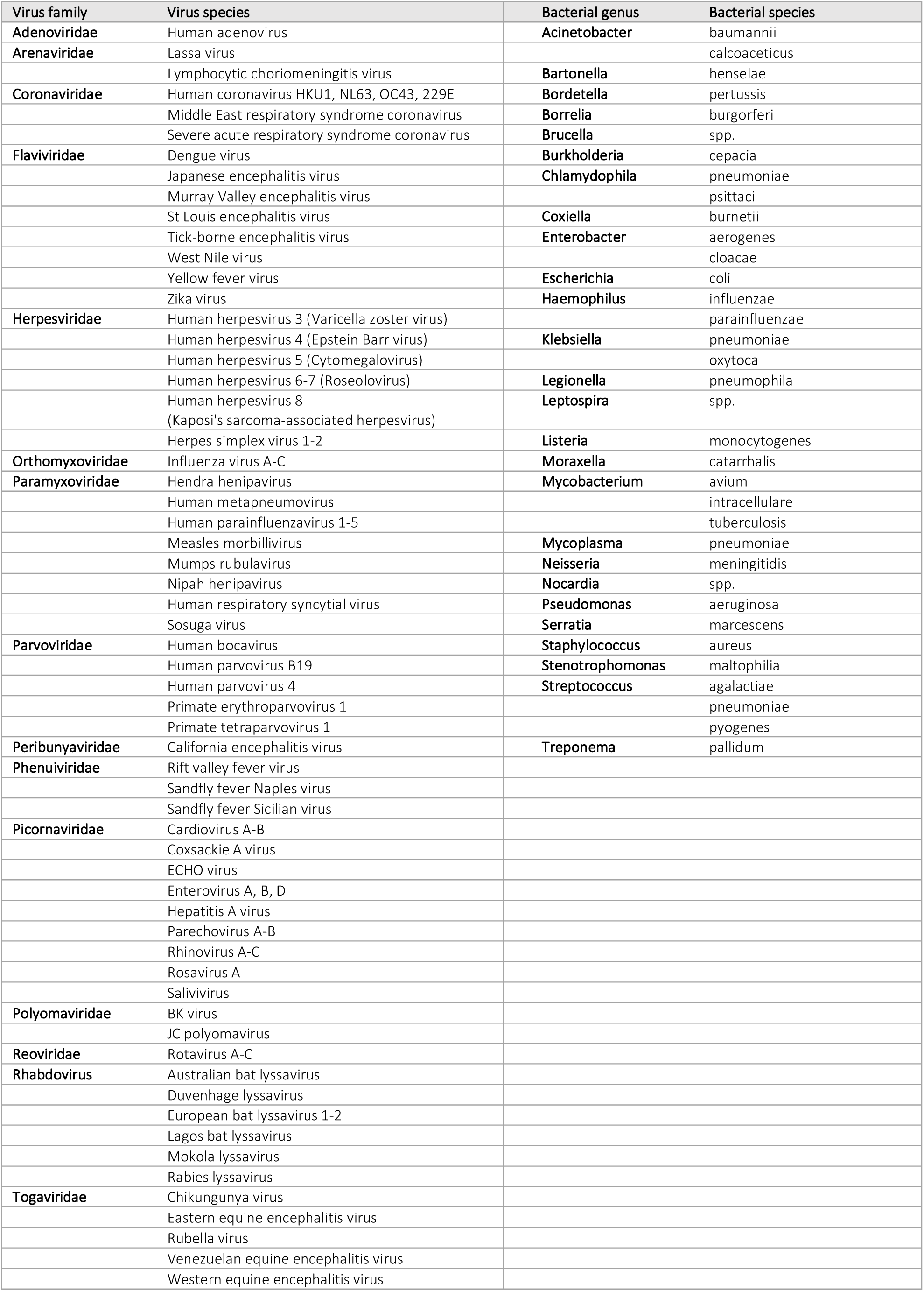
Organisms included in the *Castanet* probe set.

We targeted similar lengths of genomic sequence for each pathogen to achieve comparable assay sensitivity, optimising the breadth of pathogens we could target and avoiding bias in favour of larger genomes. For each of the viruses, we downloaded from NCBI RefSeq the full set of complete genomes available at 1^st^ August 2015. Except for each of the included herpesviruses, whose genomes exceed 100 kbp and where we chose a low-diversity region of ∼20kb, for each virus we constructed a whole-genome alignment using MAFFT^38^ from which to design probes. For bacterial species, we took advantage of the ribosomal multilocus sequence typing (rMLST) scheme, which targets 53 genes encoding ribosomal proteins present in all bacteria and resolves bacteria to a sub-species level^39^, extracting sequences from the rMLST database on 11 December 2015.

In previous work we observed that sequence capture efficiency is preserved even when probe and target sequences diverge substantially, and that exploiting sequence similarity to avoid redundancy can improve the efficiency of probe design without sacrificing performance^28^. Accordingly, for each sequence alignment we constructed a UPGMA tree using pairwise Hamming distances, within which we identified clusters such that all sequences were less than 5% divergent from one another. This process defined a set of 5.86 × 10^6^ bases of cluster consensus sequences that were used to synthesise a panel of 52,101 Agilent SureSelect RNA probes, each of 120 nucleotides on the complementary strand.

### Clinical sample collections for *Castanet* evaluation

We assembled a large collection of samples from two clinical study cohorts in order to assess the performance of our sequencing and analysis workflow.

#### The GAinS Study

(https://ukccggains.com/) is a multi-centre clinical study of sepsis conducted in the UK, including n=2055 sepsis cases for which one or more clinical samples and extensive clinical information were collected [REC: 05/MRE00/38, 08/H0505/78]. We analysed plasma samples from n=573 patients diagnosed with sepsis secondary to community-acquired pneumonia, comprising n=126 ‘known’ cases where clinical microbiology had identified a pathogen (although not necessarily from the available plasma sample) and n=447 ‘unknown’ cases chosen from the whole cohort because no pathogen had been identified in any sample from that case. Accordingly, our sepsis sample collection cannot be considered a random cross-section of cases, nor a full set of samples for each case from which a pathogen might be identified. It was therefore necessary to confirm the pathogen status of the set of sepsis plasma samples used as ‘known’ samples to validate our methodology. In addition, aggregate results from the available sample collection should not be considered to be representative of the sepsis cohort, with respect to rates of pathogen identification or pathogen frequency.

#### The ChiMES Study

(http://www.encephuk.org/studies/ukchimes) is a multi-centre clinical study of >3,000 children with suspected meningitis and encephalitis conducted in the UK, where one or more clinical samples and extensive clinical information were collected [REC: 11/EM/0442]. We obtained CSF samples from n=243 cases, divided into n=108 ‘known’ cases for which a pathogen had been detected and n=135 ‘unknown’ cases for which no pathogen had been detected in the CSF. In general, only one CSF sample had been taken per case, so we were able to assume that our ‘known’ samples contained the relevant pathogen. Since the complete sample set comprised an essentially random selection from the case cohort, we consider the rates of identification and frequencies of pathogens to be representative of UK childhood meningitis cases within the study period and locations.

Based on the clinical cohort descriptions above, we note that the ‘unknown’ sample sets are likely to have lower pathogen levels, and/or fewer samples with detectable pathogen material, and/or a higher proportion of pathogens for which effective testing has not been applied. Furthermore, the successful clinical microbiology pathogen identifications among sepsis cases come from a larger number of more diverse sample types (many of which are unavailable for this study) than those in the meningitis cases, emphasising challenges in identifying pathogens among the ‘unknown’ samples. Further details of cohorts are in the **Supplementary Information**.

### A combined library preparation method enables sequencing of both DNA and RNA from microbiological samples

We combined a sample purification method that isolates DNA and RNA together with a single library preparation workflow for both RNA and DNA (**Methods**) in order to minimise sample wastage, reduce costs and avoid bias. We used spike-ins of plasmid DNA and ERCC RNA to confirm that our library method could recover DNA and RNA sequences with similar efficiencies (**Supplementary Information**).

### *Castanet* provides sensitive, quantitative detection and sequencing of viruses and bacteria

To assess the quantitative range of detection of our method, we used dilutions of a commercially available mixture of viruses, designated Viral Multiplex Reference 11/242 (VMR) (National Institute of Biological Standards and Control, UK)^36^. From two undiluted VMR replicates, *Castanet* sequencing yielded 9.1×10^7^ and 10.9×10^7^ reads respectively (**Supplementary Information**). All 21 viruses in VMR for which we had enrichment probes were detected, with at least 8.65 reads per million in enriched samples (**Supplementary Figure 1)**. Our method, which has been designed for high-throughput processing of samples in a single assay, compared favourably with the multiple methods adopted for detection of all VMR viruses in 15 other laboratories^36^.

We compared viral load with enriched sequence yield for the subset of five VMR viruses that had been quantified by the supplier using qPCR. For each virus, the number of deduplicated *Castanet* reads increased linearly with a similar constant of proportionality to input copy number across a 3-log dilution series (**Figure 1a**). The number of unique reads obtained per virus genome copy differed between viruses, presumably because of differences in sequence target length, the efficiency of sequencing library formation and perhaps, the calibration of qPCR assays.

**Figure 1:**
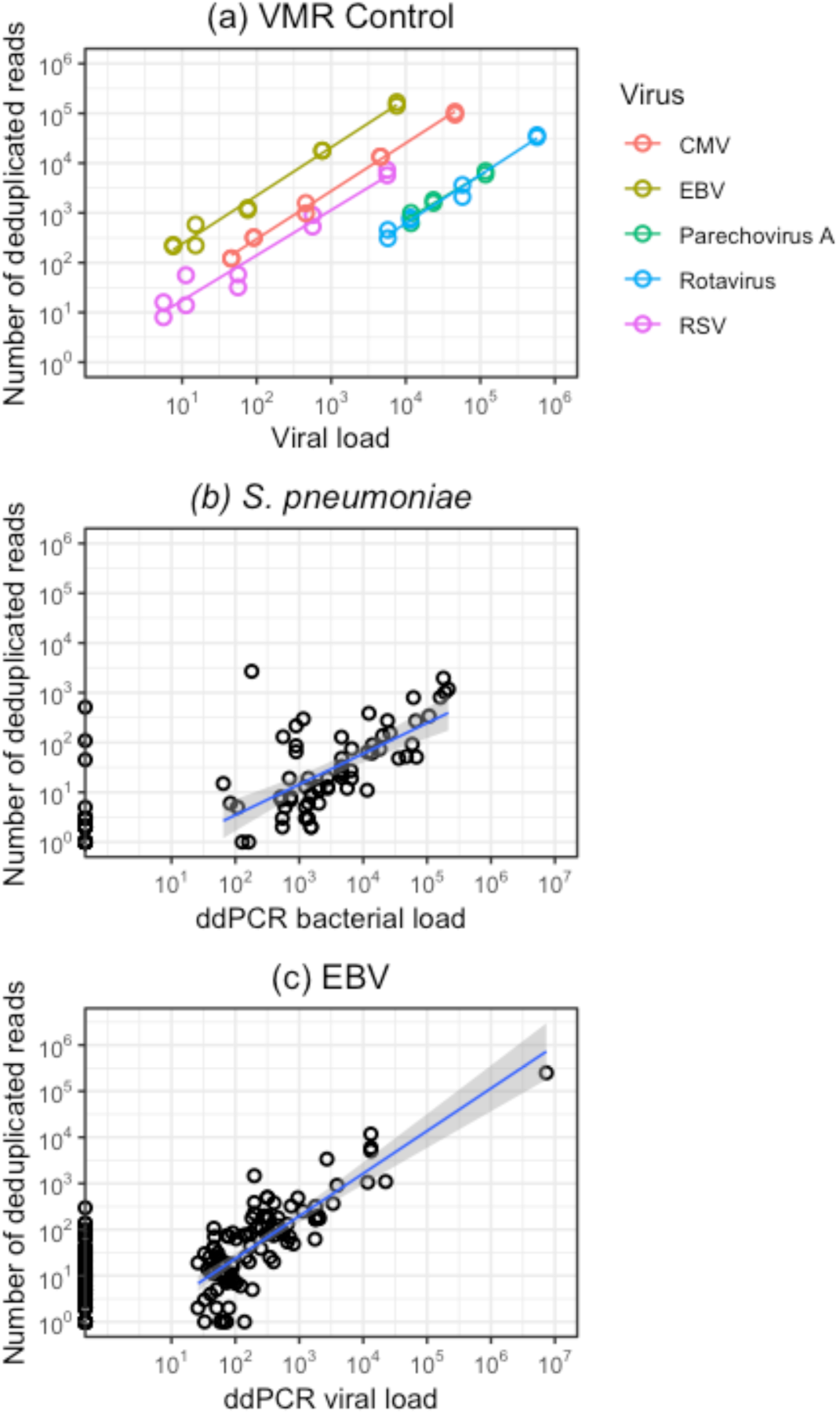
Organism load and sequencing yield in control and sepsis samples. (a) Viral Multiplex Reference reagent was prepared for sequencing at a range of dilutions (neat, 1:10, 1:100, 1:500, 1:1000) in two replicates. Deduplicated read yield was plotted against input viral load for the five viruses that had been quantified by the manufacturer using qPCR. (b) and (c) Pathogen load and sequencing yield in clinical sepsis samples. Deduplicated read yield was plotted against pathogen load, estimated by ddPCR, in samples from a subset of cases in which (b) *Streptococcus pneumoniae* (n=102) or (c) Epstein-Barr Virus (EBV) (n=199) was detected by sequencing. A linear relationship between pathogen load and sequencing yield was observed for each organism (*S. pneumoniae*: R^2^ = 0.449, p = 8.8 × 10^−5^; EBV: R^2^ = 0.702, p = 2.9 × 10^−16^). The limit of detection of the ddPCR assay was considered to be (b) 46 copies/ml, (c) 26 copies/ml. Sequencing yields for ddPCR-negative samples are shown but not included in the linear regression. VMR, Viral Multiplex Reference; CMV, human cytomegalovirus; EBV, Epstein-Barr virus; RSV, respiratory syncytial virus.

Similarly, we observed a striking linear relationship between input pathogen load and sequencing yield in a subset of clinical sepsis samples for which we had quantified bacterial and viral load of *Streptococcus pneumoniae* (*Spn*) (n=102) and Epstein-Barr virus (EBV) (n=199) by digital droplet PCR (ddPCR) (**Figure 1b, 1c**). Quantitative yield from targeted enrichment across a wide range of input concentrations is consistent with previous work^28^. In these experiments, sequence enrichment increased the yield of pathogen-specific reads by ∼10,000 times for an equivalent amount of sequencing (**Supplementary Figure 2**), implying that batches of a convenient size for larger studies, such as plates of 96 samples prepared and pooled together, would attain at-least-equivalent sensitivity with the same amount of sequencing as a single sample undergoing standard shotgun metagenomic sequencing, massively reducing assay costs.

### Large-scale analysis of clinical samples with *Castanet*

We successfully sequenced and included in our analysis a total of 854 samples, derived from 243 ChiMES meningitis cases, 27 non-meningitis negative-control CSF samples, 573 GAinS sepsis cases and 11 negative-control plasma samples. The 243 meningitis cases formed an approximately random sample of disease that comprised 122 for which a pathogen had been identified by clinical microbiology (108 from CSF only plus 14 from a blood sample plus possibly also CSF) and 121 where no pathogen had been identified before sequencing. The sepsis cases comprised 126 for which a pathogen had been identified and 447 chosen for study because no pathogen had been identified in any relevant sample.

### *Castanet* detects clinically relevant pathogens in CSF and plasma samples

In order to evaluate the ability of *Castanet* to identify pathogens in real clinical samples, we compiled a ‘truth dataset’ of samples whose status for particular pathogens was known with confidence. For meningitis cases, we accepted the pathogen identification in the case record as the truth state for microbiology-positive samples. For the 126 pathogen-positive CAP sepsis cases, the pathogen identification had often been made in a sample other than plasma and most of the plasma samples in our collection had been obtained after administration of antibiotics, two situations in which plasma levels of pathogens might have been undetectable by any method. Accordingly, we used a positive result by ddPCR for *S. pneumoniae* or EBV to define a sample subset with which to learn the characteristics of pathogen-true-positive sequencing data.

Another key issue in interpreting metagenomic sequencing data, especially in large pools of samples, is to distinguish low-level positives from true-negative samples. We therefore used the opportunity provided by the *S. pneumoniae* ddPCR assays to estimate the threshold for considering a sample sequence-positive in the current study (**Supplementary Figure 3**), deciding to exclude ddPCR-negative samples with fewer than either 72 total or 4 de-duplicated sequence reads from any single pathogen from consideration as sequencing-positive.

Combining the above criteria, we identified 100 ChiMES meningitis CSF and 107 GAinS sepsis plasma samples for inclusion as samples with known pathogen status. To these we added samples included in the cohorts as negative controls (CSF from non-meningitis patients, n=23; and plasma from sepsis-negative patients, n=17) to provide instances of pathogen-negative data. Plasma and CSF samples that were microbiology-positive for a particular pathogen were deemed negative for other pathogens. Reads aligning to viruses known to reactivate in sepsis (herpes simplex virus (HSV), cytomegalovirus (CMV), human herpes virus 6 (HHV6), JC virus) were excluded from the analysis, apart from those EBV samples where ddPCR data was available.

We randomly allocated the 247 samples defined above to training and test datasets in an 80:20 ratio. We used the training dataset (197 samples: 95 CSF; 102 plasma) to train a random forest classifier that used a set of variables derived from the sequencing data (details in **Supplementary Material**) to derive a score between 0 and 1 to indicate whether it was positive for each organism with reads in a sample. Some samples contained reads from multiple organisms and the random forest returns a score for each one of those organisms.

The test dataset comprised 50 samples (28 CSF; 22 plasma). We selected a cut-off random forest score of 0.465 for classification of the test set, to appropriately weight specificity over sensitivity (**Supplementary Figure 4**). At this threshold, there were 5 false negatives and one false positive in the ChiMES test dataset and one false negative and 3 false positives in the GAinS test dataset. In the combined set of test samples the sensitivity was 86.7% (39 of 45 true positives, Figure 2) and the specificity was 98.6% (283 of 287 true negatives). Among the most informative sequencing data metrics for prediction were the numbers of total and deduplicated reads matching a pathogen, taken as the respective proportions of reads aligning to all pathogens in the probeset, and whether a high proportion of the targeted region (regions in our probeset) for that pathogen were covered by reads (**Supplementary Figure 5)**. In meningitis predicted-positive samples a median 91% of the genomic target sequence for the implicated pathogen was covered by at least 2 reads. Since excluding pathogens from a diagnosis can also be clinically useful, we assessed the performance of the method in predicting negative status. With a random forest score threshold of 0.015, we predicted 59.2 % of true negatives with a specificity of 97.8%, implying that for many samples it is possible to exclude many possible pathogens without erroneously ruling out true positives.

**Figure 2:**
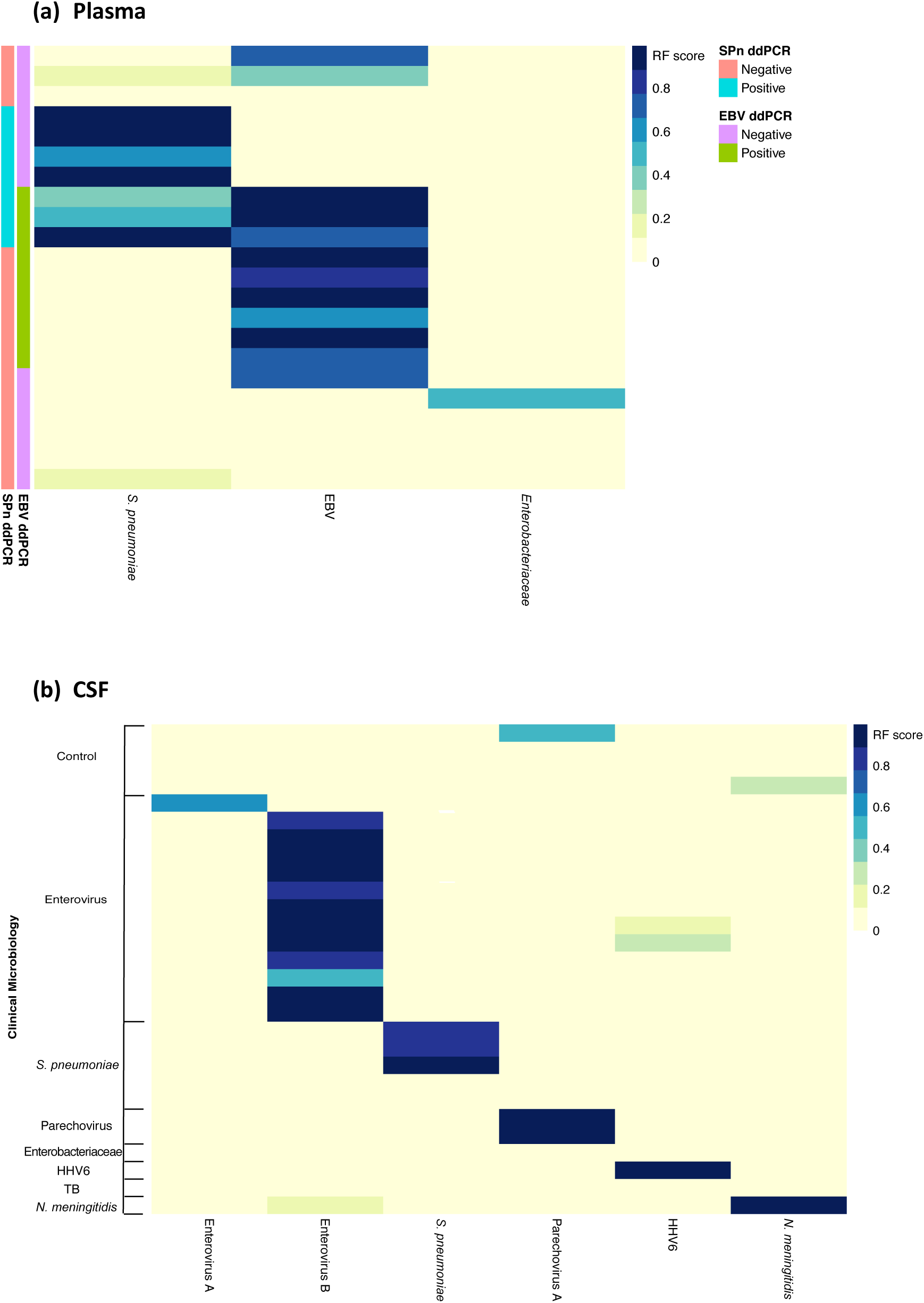
Performance of *Castanet* in clinical samples with a positive microbiology diagnosis. The test dataset included 50 samples: (a) 22 plasma samples (20 GAinS, 2 sepsis-negative controls); (b) 28 CSF samples (24 ChiMES, 4 meningitis-negative controls). The combined overall test dataset specificity and sensitivity was 0.986 and 0.867 respectively. [ChiMES=Childhood Meningitis and Encephalitis Study. GAinS=Genomic Advances in Sepsis. RF=Random Forest. EBV=Epstein-Barr Virus. SPn=Streptococcus pneumoniae. HHV6=human herpes virus 6. ddPCR=droplet digital PCR]. Only organisms detected in each sample set are included as columns in each panel.

### *Castanet* detects pathogens in samples from patients with no previous pathogen identification

We analysed the remaining 121 ChiMES meningitis CSF and 447 GAinS sepsis plasma samples, in which no causative pathogen had been identified by routine clinical microbiology. *Castanet* identified one or more pathogens in over one third of samples in both meningitis (41 samples, 34%) and sepsis (165 samples, 37%), including both bacteria and viruses that in many cases were likely to have been causative (**Figure 3**). Among such pathogens, instances of EBV, HHV6, HSV, JC virus and HCMV in sepsis may represent viral reactivation in the context of critical illness, while *Burkholderia cepacia* and *Nocardia asteroides* sequences most probably represent contamination of samples^40^. Excluding these likely reactivations and contaminants, *Castanet* made 39 new detections of clinically relevant pathogens in meningitis patients with negative clinical microbiology, and 50 such new detections in sepsis patients, comprising 32% and 11% of previously unresolved cases respectively (**Table 2** and **Table 3**).

**Figure 3:**
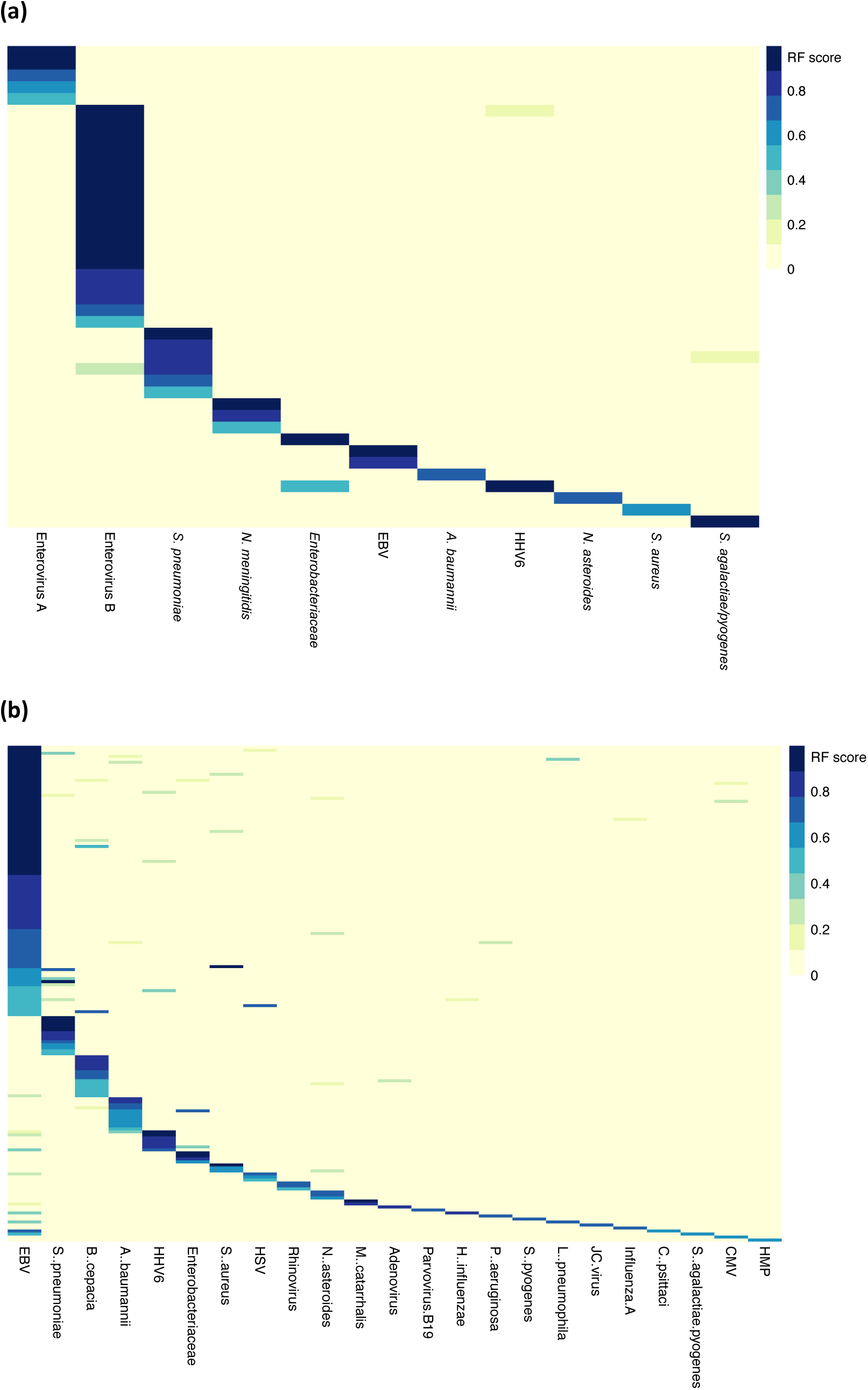
Performance of *Castanet* in samples with no clinical microbiology diagnosis. Panels show as rows (a) 41/121 ChiMES samples; (b) 165/447 GAinS samples; for samples with at least one organism detected at a random forest score RF >0.465. Only organisms detected in each sample set are included as columns in each panel.

**Table 2:**
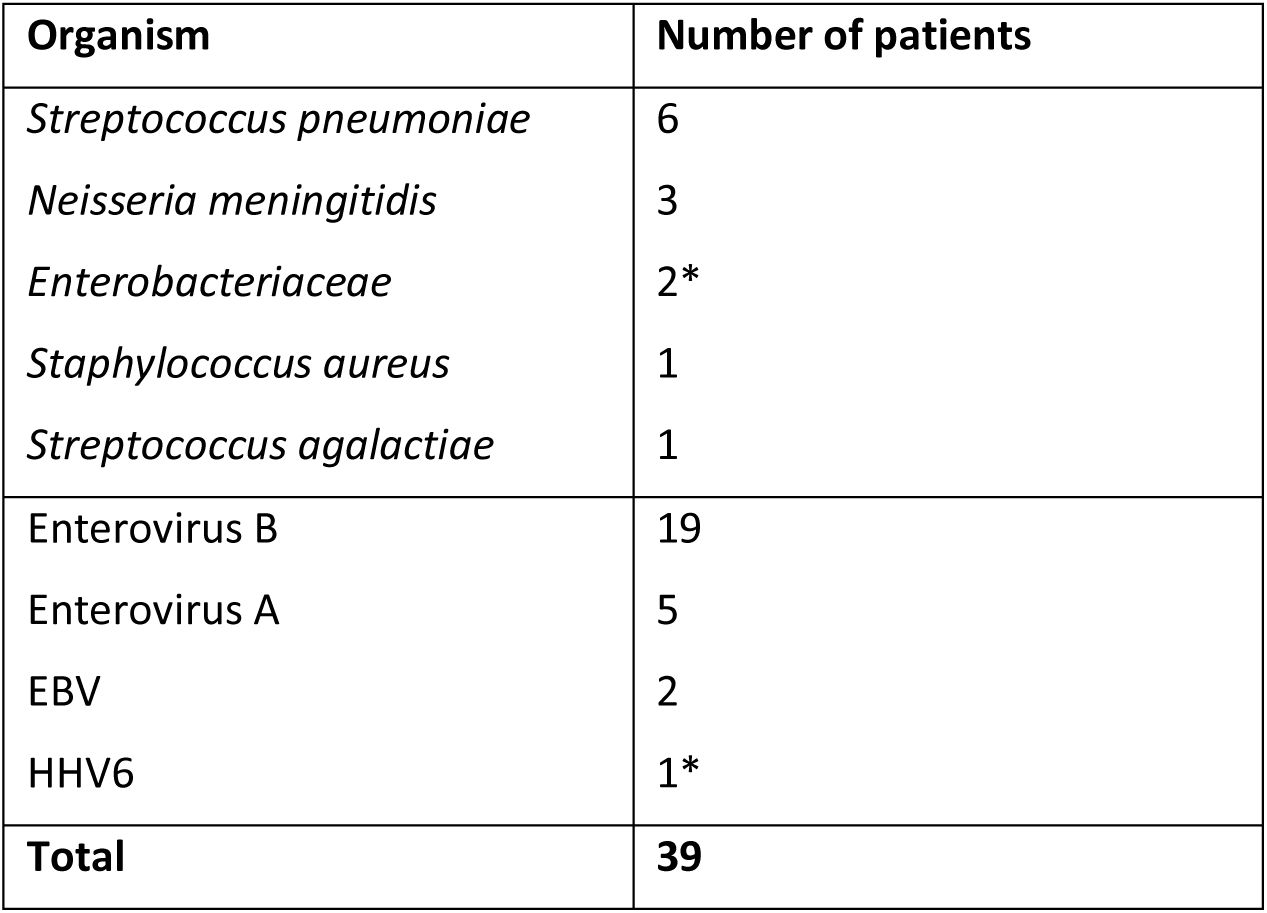
New pathogen identifications made by *Castanet* among meningitis CSF samples.

**Table 3:** New pathogen identifications made by *Castanet* among sepsis plasma samples.

| Organism | Number of patients |
| --- | --- |
| *Streptococcus pneumoniae* | 16 |
| *Acinetobacter baumannii* | 11 |
| *Enterobacteriaceae* | 5 |
| *Staphylococcus aureus* | 4 |
| *Moraxella catarrhalis* | 2 |
| *Haemophilus influenzae* | 1 |
| *Pseudomonas aeruginosa* | 1 |
| *Streptococcus pyogenes* | 1 |
| *Legionella pneumophila* | 1 |
| *Chlamydia psittaci* | 1 |
| Rhinovirus | 3 |
| Adenovirus | 1 |
| Parvovirus B19 | 1 |
| Influenza A | 1 |
| Human metapneumovirus | 1 |
| **Total** | **50** |

In the complete meningitis cohort, *Castanet* improved the overall rate of pathogen identification from 50% (122/243) to 69% (175/243) compared with clinical microbiology alone. Specifically, in CSF samples, *Castanet* identified a likely causative pathogen in 149 of 243 cases (61%) compared with 108/243 (44%) for clinical microbiology (**Table 4**). **Table 5** gives equivalent totals for the sepsis plasma sample set.

**Table 4.** Pathogens identified in all meningitis clinical cases / meningitis CSF samples.

| Organism | Clinical microbiology: CSF only | Clinical microbiology: CSF + blood | Clinical microbiology + *Castanet* | Clinical microbiology + *Castanet* | *Castanet* | Prevalence in sample set |
| --- | --- | --- | --- | --- | --- | --- |
| Unknown | 135 (56%) | 121 (50%) | 80 | 80 (33%) | 94 (37%) | 42% |
| Enterovirus | 45 (19%) | 45 (19%) | 75 | 163 (67%) | 74 (30%) | 31% |
| Human parechovirus | 14 (6%) | 14 (6%) | 15 |  | 13 (5%) | 4% |
| *Streptococcus pneumoniae* | 11 (5%) | 15 (6%) | 24 |  | 18 (7%) | 5% |
| *Neisseria meningitidis* | 8 (3%) | 13 (5%) | 17 |  | 15 (6%) | 6% |
| Other | 30 (14%) | 35 (15%) | 44 |  | 29 (12%) | 11% |
| **Total** | **243** | **243** | **255 \*** | **243** | **243** |  |

\* 12 samples called differently by lab and *Castanet* appear in two categories in the column above.

**Table 5.** Pathogens identified in all sepsis clinical cases / sepsis plasma samples.

| Organism | Clinical microbiology | Clinical microbiology + *Castanet* | *Castanet* | Prevalence in sample set |
| --- | --- | --- | --- | --- |
| Unknown | 447 (78%) | 397 (69%) | 488 (85%) | 60% |
| *Streptococcus pneumoniae* | 92 (16%) | 108 (19%) | 45 (8%) | 15% |
| Other bacterial | 6 (1%) | 33 (6%) | 31 (5%) | 18% |
| Influenza | 13 (2%) | 14 (2%) | 1 (0%) | 5% |
| Other viral | 15 (3%) | 21 (4%) | 8 (1%) | 3% |
| **Total** | **573** | **573** | **573** |  |

In several cases sequencing suggested results that differed from the recorded clinical microbiology. In some instances, sample-specific reads corresponding to the microbiological diagnosis remained below the threshold of the detection model. Many discrepancies were difficult to resolve (for example where a different pathogen may have actually been present in another sample from the same case), so we focused on Enterovirus detections, where there was good information on PCR-based testing, at least at the case level (**Table 6**). There are 18 samples which are PCR-negative, but *Castanet*-positive.

**Table 6.** Resolving Enterovirus identifications in the ChiMES meningitis CSF sample set.

|  | Clinical microbiology status |  |  |  |
| --- | --- | --- | --- | --- |
|  | PCR-positive | PCR-negative | Not tested | Total |
| *Castanet*-positive | 44 | 18 | 16 | **78** |
| *Castanet*-negative | 1 | 142 | 50 | **193** |
| **Total** | **45** | **160** | **66** | **271** |

Our investigation provided a simple example of one potential application of the methodology: more extensive, comprehensive testing for a wider panel of potential pathogens can lead us to shift our understanding of (i) the complement and (ii) the frequency distribution of pathogens causing a disease in a specific population. In the meningitis cohort we detected (and in most cases derived a complete genome sequence for) a substantial number of diverse enteroviruses, such that the estimated frequency of enteroviruses among all identified pathogens (74 Enterovirus, 75 other pathogens) in the sample defined using *Castanet* was significantly higher than that defined using clinical microbiology methods alone (45 Enterovirus, 77 other pathogens; p = 0.03, Fisher’s Exact Test).

## Discussion

Our workflow, *Castanet*, is a versatile probe-based enrichment sequencing method that addresses several of the issues associated with sequencing for pathogen detection. Our method is sensitive and cost-effective, combining the analysis of RNA and DNA from the same material in a single protocol and enriching for pathogens of interest using a modestly sized panel of probes that efficiently targets a diverse range of viral and bacterial pathogens and can be tailored to particular settings.

*Castanet* attains molecular sensitivity comparable to PCR with a yield of pathogen sequence reads that is proportional to pathogen load. In this study we have adopted a probe panel targeting members of 17 virus families and 35 bacterial species relevant to two disease presentations. With approximately 10,000-times enrichment of targeted sequences, the method can analyse many samples in parallel using the same amount of sequencing that could otherwise be needed for one sample.

We evaluated the *Castanet* methodology on a large sample set from two densely phenotyped disease cohorts collected across multiple centres in the UK. We selected these cohorts as the most challenging situations on which an assay like *Castanet* would be applied in a diagnostic laboratory setting. In both sepsis and meningitis, the absolute rate of identification of pathogens tends to be low despite extensive laboratory testing, and many samples may be taken after administration of antibiotics. In the setting of sepsis secondary to pneumonia, plasma may also contain little or no pathogen material from agents such as influenza virus A that had been responsible for the initial infection.

In spite of difficulties in describing a ground-truth dataset of known positives and known negatives, we clearly demonstrate that *Castanet* embodies high sensitivity and specificity in a single test. Acknowledging the complexities of interpreting metagenomics data, we show that the characteristics of pathogen-positive and pathogen-negative samples can be defined using a random forest model that combines data across pathogens and diseases.

A key aim of our study was to evaluate the performance of the *Castanet* method in increasing the number of pathogen identifications made from challenging samples. We identify a substantial number of previously unrecognised infections and show that *Castanet* can be used to update estimates of population pathogen frequencies derived from conventional testing.

The *Castanet* method has a variety of prospective applications. A key difference from other targeted assays is that there is no need to decide for each sample what tests to perform; with probe-enrichment the user can in principle (and in practice) construct a very long list of plausible pathogens to test for with equivalent sensitivity. By providing a wide specificity in a single test on large batches of samples, *Castanet* will be useful in investigating the performance and coverage of existing testing regimens and platforms (for example to check whether a test covers all circulating variants or the set of tests in use includes all common pathogens in the population). Similarly, *Castanet* could define with minimal bias the distribution of pathogens associated with a disease in a particular population, particularly for low- and middle-income countries where treatment may be based on untested assumptions. In principle, probe enrichment sequencing could allow clinicians for the first time to exclude many possible causes of infection at the same time, in contrast to current methodologies.

In this study, we provide an example of a further use of the *Castanet* workflow by deriving complete genomes of diverse viruses on a large scale, with sensitivity comparable to PCR but with greater robustness and without the requirement for assay optimisation. In addition to the diverse set of Enterovirus genomes generated within this study we have started using *Castanet* as a high-throughput pipeline for sequencing viruses from nasal swab samples in acute respiratory infection (manuscript in preparation).

Future work is envisaged to achieve better resolution of pathogenic species within clusters of closely related bacteria such as the Gram-negatives, Streptococci and Staphylococci, including by focusing on bacterial sequences of particular clinical interest such as factors associated with virulence and anti-microbial resistance. Further challenges posed by the need to optimise sensitivity by improving the efficiency with which pathogen nucleic acids are processed into sequencing libraries that could also incorporate unique molecular identifiers to enhance subsequent data analysis.

Metagenomics using short read sequencing has long promised a new approach to microbiological testing, with limited impact as a patient-focused test because of unwieldy preparation methods and slow, costly and high-volume sequencing runs. On the other hand, streamlined sample-processing and immediate dataflows mean that nanopore-based sequencing is already close to providing a workflow compatible with a fast-turnaround lab-based or point-of-care test, even if data volumes are not yet sufficient for sensitive metagenomics at a reasonable cost. Probe enrichment can provide benefits for both approaches, enabling high-throughput batch testing and, if capture times can be reduced, enhancing the sensitivity and therefore speed and cost of nanopore sequencing. In either form, our approach could be useful in routine microbiology as a backup to fast-turnaround tests or to exclude large numbers of rare pathogens with turnaround of results in a few days.

*Castanet* and related targeted metagenomics methodologies provide a significant step towards the widespread use of metagenomics in identifying and characterising pathogen sequences in large-scale clinical studies. By providing a single workflow, *Castanet* promises to simplify and generally enable phylo-epidemiological studies on many viruses and to be valuable in defining causes of infection at the population level, with benefits for the quality of microbiological testing.

## Methods

### Experimental procedures

#### Nucleic acid extraction

Total nucleic acids were extracted using the NucliSENS easyMag platform (Biomerieux) from up to 500ul of plasma or CSF where available and the extracted nucleic acids were eluted in 25ul of kit elution buffer and stored at −80°C.

#### Library preparation

RNA was reverse-transcribed with random priming using the NEBNext Ultra Directional RNA Library Prep Kit for Illumina (New England Biolabs) with 5ul sample input. We made modifications to the manufacturer’s guidelines, including fragmentation for 4 minutes at 94°C and omission of Actinomycin D at first-strand reverse transcription, and we stopped the protocol after generation of double-stranded cDNA. The resulting mixture of cDNA and sample DNA (normalised to 5ng/ul or the maximum available) was processed using the Nextera DNA Library Preparation Kit (Illumina) according to the manufacturer’s guidelines, including library amplification for 15 PCR cycles using custom indexed primers and post-PCR clean-up with 0.85x volume Ampure XP (Beckman Coulter). Libraries were quantified using Quant-iT PicoGreen dsDNA Assay Kit (Invitrogen) and analysed using Agilent TapeStation with D1K High Sensitivity Kit (Agilent) for equimolar pooling.

#### Probe-based enrichment

1ug of each indexed pooled library was enriched using the Agilent SureSelect^*XT*^ Target Enrichment System for Illumina Paired-End Multiplexed Sequencing Library protocol with one major modification to the recommended protocol: the capture was performed on the post-PCR indexed pool, using oligonucleotide blockers complementary to adapter sequences.

#### Sequencing

Sequencing was performed on the Illumina MiSeq or HiSeq 4000 platforms generating either 75-bp or 150-bp paired-end reads.

#### Digital Droplet PCR (ddPCR)

Aliquots of extracted nucleic acid from plasma were processed in triplicate (1.5ul per replicate) following the recommended workflow (WX200 ddPCR system, Bio-Rad). Custom PrimeTime (IDT) primer/probe sets targeting *S*. pneumoniae,^41^ and Epstein-Barr Virus (EBV),^42^ were designed based on published sequence data. Full details are included in the Supplementary Material.

### Bioinformatics and statistical analysis

#### Sepsis and meningitis samples and uninfected controls

De-multiplexed sequence read-pairs were trimmed of adapter sequences using Trimmomatic v0.36, with the ILLUMINACLIP options set to “2:10:7:1:true MINLEN:80”, using the set of Illumina adapters supplied with the software.^43^ The trimmed reads were then classified using Kraken v1^44^ using a custom database containing the human genome (GRCh38 build), all RefSeq bacterial and viral genomes, and a selection of fungal genomes that were most likely to be associated with cases of meningitis^45^. These were: *Aspergillus fumigatus, Candida* spp., Coccidioides spp., *Cryptococcus* spp., *Histoplasma capsulatum, Paracoccidioides brasiliensis*, and *Pneumocystis* spp. Reads identified as bacterial or viral and unclassified reads were aligned using BWA v0.7.12^46^ with default settings to a multi-fasta reference of consensus sequences corresponding to the enrichment probe targets, augmented with sequences of known or suspected contaminants. These included (i) reagent contaminants (Alteromonas and Achromobacter spp.), (ii) genomes of two viruses known to have been sequenced on the same flow cell: MVMPCG and Echovirus 7; and (iii) the rMLST sequences of commensal *Streptococcus* species species that were thought to be likely contaminants in clinical samples (Supplementary data file).

#### Duplicate reassignment

We first corrected our sequencing results for index misassignment, a well-recognised issue with multiplexed sequencing where a small proportion of reads in each sample represents an incorrectly identified read that had come from a nearby optical cluster. For each sequencing pool, we identified PCR-duplicated reads and reassigned all reads in each duplicate cluster to the sample with the highest number of reads in that cluster.

#### Collection of sequencing statistics

Following duplicate reassignment, for each sample and target organism we calculated a set of descriptive statistics, which included sequencing depth with and without deduplication, and coverage of target sequences at a various depth thresholds. The collected statistics were combined with available laboratory data on sample positivity, and with ddPCR results where these were available. The resulting data frame was used to train a Random Forest model.

#### Random Forest classification model

The function randomForest from the R package of the same name was used to derive an algorithm to classify reads aligning to each organism in a sample as positive or negative (details in **Supplementary Information**).

## Study Information

### Contributions

C.G., T.G., A.A., A.T., D.B., P.P., E.B., A.P., M.S., J.K., and R.B. conceived and designed the study. C.H. and P.H. (GAinS), and M.S., N.M., S.D., M.G., A.P., and T.S. (ChiMES) were responsible for clinical study setup, patient recruitment and sample collection. A.A. designed the probe set. C.G., I.E., J.B., M.M., M.S. and R.B. contributed to probe design. C.G., M.C., A.T., D.B., P.P., and R.B. contributed to laboratory methods development. A.B. and C.G. undertook the nucleic acid extractions. M.C., A.T., H.S., and C.G. undertook the library preparations. C.G., T.G., R.B. and A.A. analysed sequence data. T.G. designed and developed the computational tools. J.K., A.P. and M.S. contributed to data analysis and interpretation. C.G., T.G., A.A., and R.B. wrote the paper. All authors read and approved the final manuscript.

### Research Ethics

#### ChiMES

part of the larger ENCEPH UK study - NRES Committee East Midlands - Nottingham 1 - 11/EM/0442.

#### GainS

REC: 05/MRE00/38 (Scotland A Research Ethics Committee), 08/H0505/78 (Berkshire Research Ethics Committee).

### Funding

This work was supported by a grant from Meningitis Research UK and by the Wellcome Trust via the Oxford-Wellcome Institutional Strategic Support Fund and core funding to the Wellcome Centre for Human Genetics (awards 090532/Z/09/Z and 203141/Z/16/Z). C.G. was supported by a Medical Research Council/British Journal of Anaesthesia/Royal College of Anaesthetists Fellowship (MR/M018725/1). M.A. was funded by the Oxford Martin School for part of this project. J.K. was supported by a Wellcome Trust Investigator Award (204969/Z/16/Z) and the NIHR Oxford Biomedical Research Centre. E.B. was funded by the Medical Research Council UK and the Oxford NIHR Biomedical Research Centre and is an NIHR Senior Investigator. Computation used the Oxford Biomedical Research Computing (BMRC) facility, a joint development between the Wellcome Centre for Human Genetics and the Big Data Institute supported by Health Data Research UK and the NIHR Oxford Biomedical Research Centre. Financial support was provided by the Wellcome Trust Core Award Grant Number 203141/Z/16/Z.

The views expressed in this report are those of the author(s) and not necessarily those of the NHS, the NIHR or the Department of Health.

## Acknowledgements

We thank Annabel Coxon, Gretchen Meddaugh and Rebecca Beckley for research support.

