## Supplementary Information for "Targeted metagenomic sequencing enhances the identification of pathogens associated with acute infection"

Goh et al.

#### Supplementary Methods

##### Clinical samples

Sepsis Adult patients admitted to intensive care with sepsis from community-acquired pneumonia (CAP) were recruited through the UK Genomic Advances in Sepsis (GAinS) study (http://www.ukccg-gains.org) (REC reference numbers: 05/MRE00/38, 08/H0505/78) from 30 participating ICUs between 2005-2015. Plasma samples (n=757) from adult patients (n=573) were collected on days 1 and/or 3 and/or 5 of ICU admission. Patients with severe sepsis (1992 ACCP/SCCM consensus definition)[1] and CAP (febrile illness associated with cough, sputum production, breathlessness, leucocytosis, and radiological features of pneumonia acquired prior to or within 2 days of hospital admission)[2] were included.

Sepsis negative Negative controls for the sepsis cohort (n=11) were recruited through the Structural Genetic Regulation of Changes in Gene Expression in Patients Undergoing Cardiopulmonary Bypass study (REC reference number: 08/H0706/94). Plasma samples were obtained prior to induction of general anaesthesia.

Meningitis Children under 16 years of age with suspected meningitis and encephalitis were prospectively recruited through the UK Childhood Meningitis and Encephalitis Study (UK-ChiMES) (http://www.encephuk.org/studies/ukchimes) (REC reference number: 11/EM/0442). Cerebrospinal fluid samples (n=254) from 244 patients were obtained by lumbar puncture at the first available timepoint.

Meningitis negative CSF from individuals requiring a lumbar puncture for reasons other than meningitis (n=22) formed a negative control group.

Process control samples Hepatitis C virus-positive plasma samples obtained through the NIHR Oxford Biomedical Research Centre Prospective Cohort Study in Hepatitis C [REC] were used in spike-in and other control experiments.

##### Digital droplet PCR

Following nucleic acid extraction, samples were processed in triplicate (1.5ul per replicate) according to the recommended workflow (QX100 ddPCR system, Bio-Rad) using the ddPCR SuperMix for Probes (Bio-Rad). Custom-designed PrimeTime (IDT) primer/probe sets were designed based on published sequence data[3 4] (**Supplementary Table 1**). PCR conditions were as follows: 95°C for 10 minutes; 40 cycles of 94°C for 30 seconds, 60°C for 1 minute; 98°C for 10 minutes and hold at 4°C. Results were analysed using the QuantaSoft software package (Bio-Rad) and samples with more than one positive droplet across the three replicates were deemed positive.

**Supplementary Table 1**: Digital droplet PCR targets and primer/probe sequences

| Organism | Target | Primer/probe | Nucleotide sequence |
|---|---|---|---|
| *Streptococcus pneumoniae* | *cpsA* 67bp amplicon | PrimeTime probe<br>Forward primer<br>Reverse primer | 5`-/5HEX/AATGTTACG/ZEN/CAACTGACGAG/3IABkFQ/-3`<br>5`-GCTGTTTTAGCAGATAGTGAGATCGA-3`<br>5`-TCCCAGTCGGTGCTGTCA -3` |
| Epstein-Barr virus | *EBNA1* 78bp amplicon | PrimeTime probe<br>Forward primer<br>Reverse primer | 5'-/56-FAM/AGGGAGACA/ZEN/CATCTGGACCAGAAGGC/3IABkFQ/-3'<br>5'-TACAGGACCTGGAAATGGCC-3'<br>5'-TCTTTGAGGTCCACTGCCG-3' |

**Positive Control: Viral Multiplex Reference 11/242**

We evaluated our workflow using a viral multiplex reference set (11/242) available through the UK National Institute for Biological Standards and Control (NIBSC)[5]. The reagent contains 25 infectious viruses covering a range of genome types (dsDNA, dsRNA, ssRNA+, ssRNA-), sizes (6.8-233.7 kb), envelope types and pre-assayed concentrations. We combined four 1ml aliquots of reagent and made two replicates of a series of five dilutions (neat, 1:10, 1:100, 1:500, 1:1000) in phosphate-buffered saline solution, forming 500ul aliquots for extraction and library preparation.

#### Supplementary Results

**Patient cohorts**

Summary clinical characteristics are summarised below for both the sepsis and meningitis cohorts.

**Supplementary Table 2:** GAinS (Sepsis) Cohort summary clinical characteristics

| Characteristic | Sepsis cohort (n=573) |
|---|---|
| Mean age (years) | 61 |
| Male sex | 324 (57%) |
| Mortality (28 day) | 121 (21%) |
| Mean duration from hospital to ICU admission | 0.8 |
| Mean Day 1 SOFA score | 6.2 |
| Mechanical ventilation | 460 (80%) |
| Vasopressors | 297 (52%) |
| Earliest ICU day of sampling | Day 1: 302 (53%); Day 3: 192 (34%); Day 5: 79 (14%) |

**ChiMES (Meningitis) Cohort**

**Supplementary Table 3:** ChiMES (Meningitis) cohort summary clinical characteristics (*to be completed*).

**Library preparation methods development**

We demonstrated that DNA and RNA are sequenced at equivalent efficiency using a combined library method. To generate a DNA sequencing control, we performed multiple restriction enzyme digest of three synthetic plasmids (3.3-6.8 kbp) resulting in 11 fragments between 379-3190 bp. We used Ambion External RNA Controls Consortium (ERCC) RNA Spike-In Mix 1 (Thermo Fisher Scientific) consisting of 92 synthetic transcripts 250-2000 nt as a sequence-specific RNA control.[6]

We isolated total nucleic acids from 5 control plasma samples using the Biomérieux Nuclisens EasyMag method. We assayed the absolute concentrations of DNA and RNA in the samples using an Agilent 2100 Bioanalyzer and added plasmid DNA and ERCC (External RNA Controls Consortium) RNA spike-in controls at 3% and 1% of the respective DNA and RNA concentrations by mass. We prepared libraries for pooling and sequencing and assessed the ratio of read counts derived from the DNA and RNA spike-ins, corrected for the 3:1 relative spike-in ratio. As intended, the DNA:RNA reads ratio varied approximately with the DNA:RNA ratio in each sample (**Supplementary Table 4**), indicating that DNA and RNA were sequenced at comparable efficiencies.

**Supplementary Table 4:** Estimated input and output DNA:RNA ratios*

| Sample | HCV-1 | HCV-2 | HCV-3 | HCV-4 | HCV-5 |
|---|---|---|---|---|---|
| Input (DNA:RNA estimated ratio) | 1.5 | 5.9 | 1.4 | 1.8 | 1.8 |
| Output (DNA:RNA total reads | 1.1 | 4.2 | 1.3 | 1.3 | 1.2 |

***** Corrected for 3:1 spike-in mass ratio).

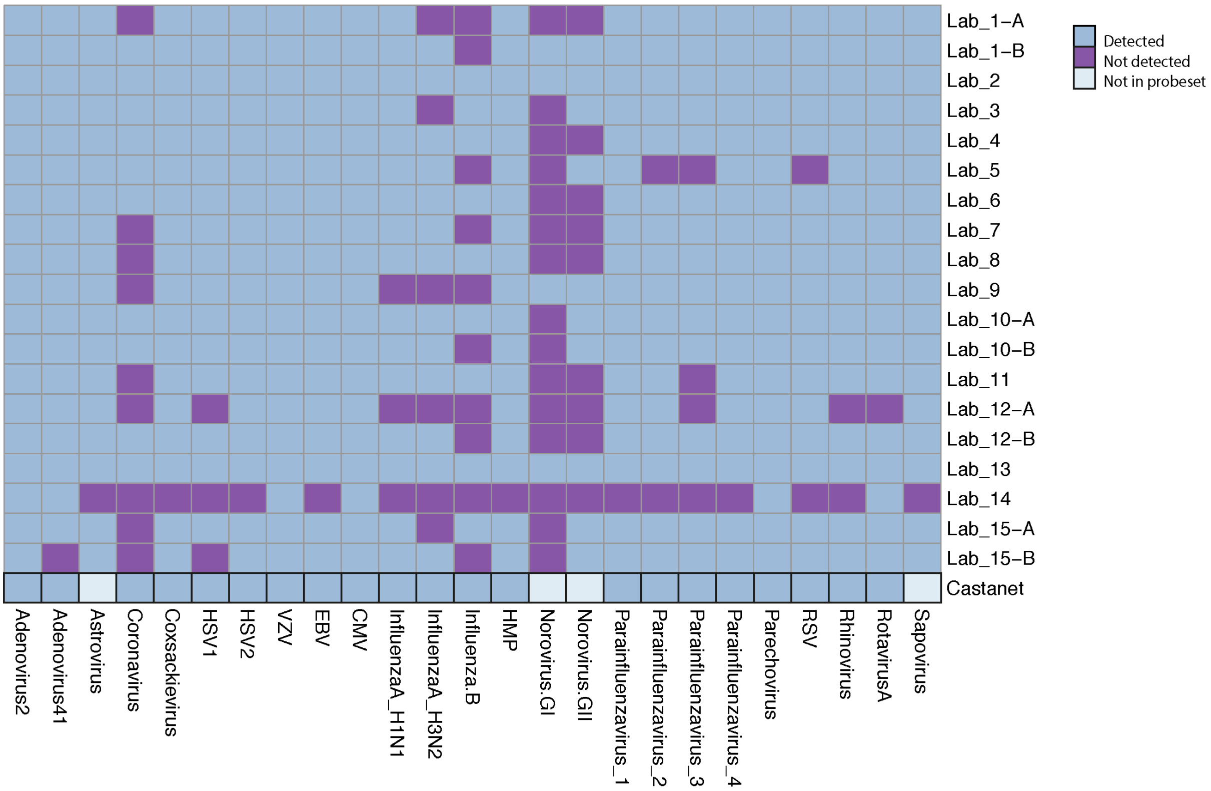


**Supplementary Figure 1.** Performance of *Castanet* in a reference set of viruses. Mee and colleagues[5] document the performance of 15 laboratories in detecting the 25 viruses in the NIBSC Viral Multiplex Reagent (VMR) 11/242 reference set. *Castanet* detected all 21 of the 25 viruses that were targeted by our probe set in an average of 91 million reads per sample. Virus abbreviations are as given in [5].

**Enrichment performance**

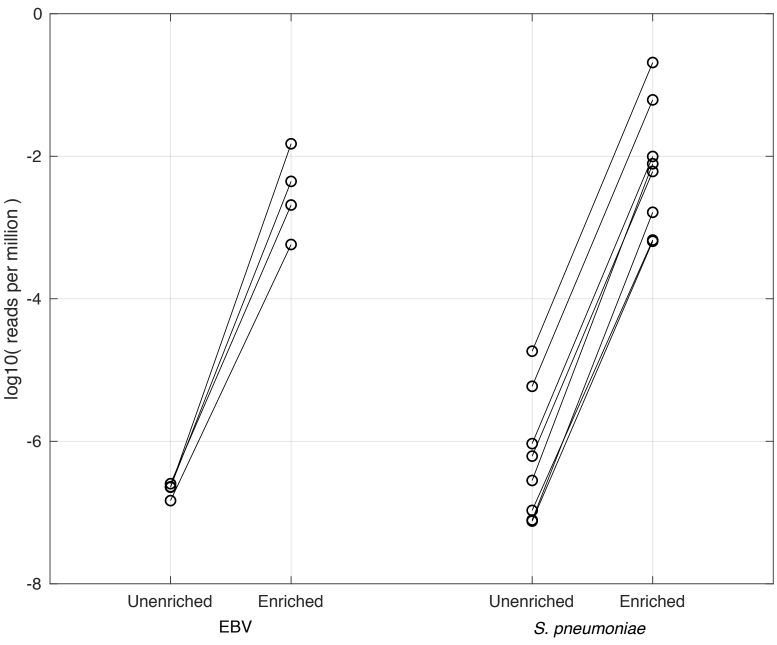


**Supplementary Figure 2.** Probe-based enrichment of a pool of libraries containing 4 EBV-positive samples and 7 S. pneumoniae-positive samples using the *Castanet* probe set. The proportion of on-target reads increases by ~10,000 times compared with libraries sequenced without enrichment. Lines link the respective samples before and after enrichment.

**Streptococcus pneumoniae limit of detection**

S. pneumoniae data from GAinS sepsis plasma samples (n=166) were used to explore the expected background level of reads at the limit of detection of our gold-standard methodology, ddPCR.

**Supplementary Figure 3**

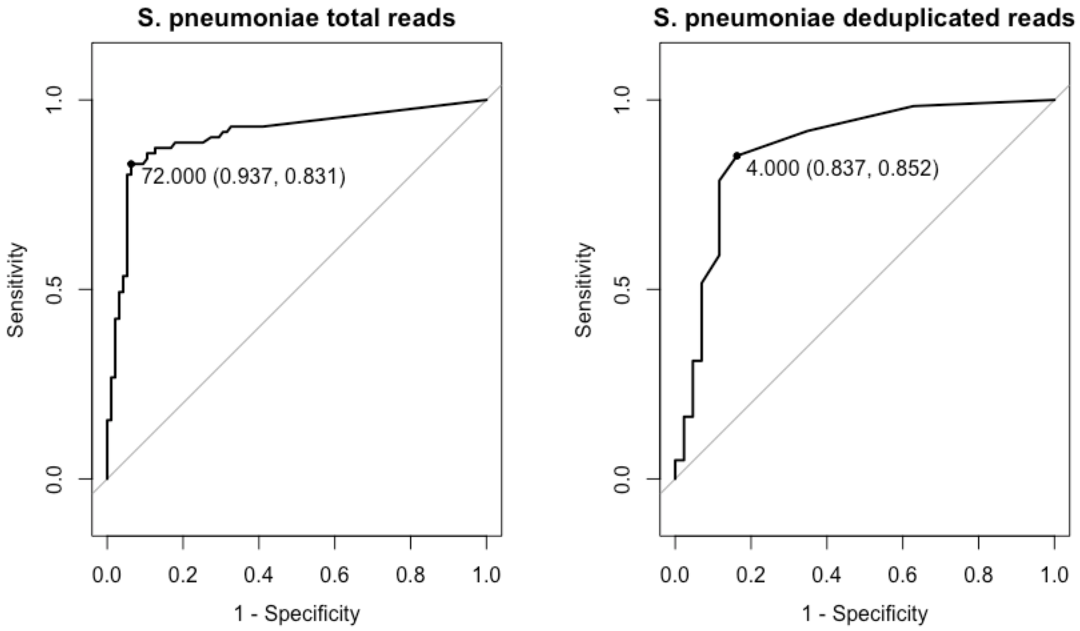


**Supplementary Figure 3**: Assessing sample read statistics as predictors of ddPCR-positive status for S pneumoniae detection. ROC curves of (a) total reads and (b) deduplicated reads as predictors. Youden's J statistic was used to select the respective thresholds of 72 total reads (specificity 0.94, sensitivity 0.83) and 4 deduplicated reads (specificity 0.84, sensitivity 0.85).

**Random forest training and calibration**

We selected 1130 observations from 247 samples (100 ChiMES meningitis CSF; 107 GAinS sepsis plasma; 23 meningitis-negative CSF; 17 sepsis-negative plasma) that were seen above the sequencing limit of detection defined above. The samples were divided into training and test datasets in an 80:20 ratio. The target for the random forest was positivity for the organism of interest, based on microbiological diagnosis (CSF) or ddPCR (plasma). We excluded samples from the training/test set if *Castanet* had identified an alternative/additional organism with a proportion of >40% total microbial reads.

**Supplementary Figure 4**

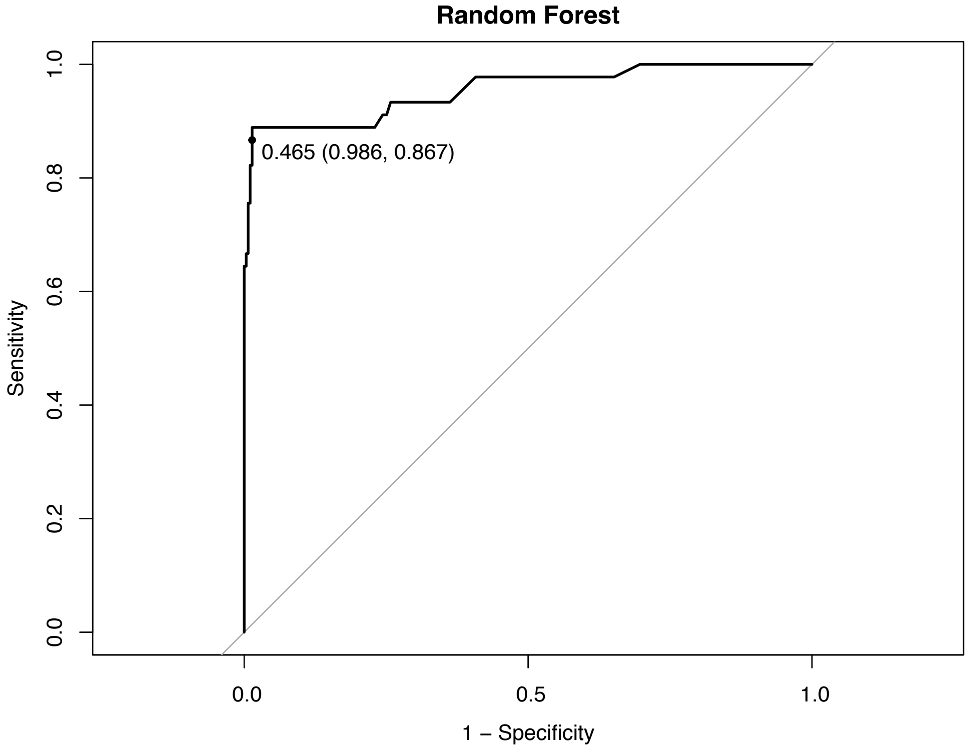


**Supplementary Figure 4**: ROC curve to choose random forest probability threshold for predicting positive samples. A threshold of RF = 0.465 was selected to call samples as positive for a particular pathogen.

### Data examples

## (a)

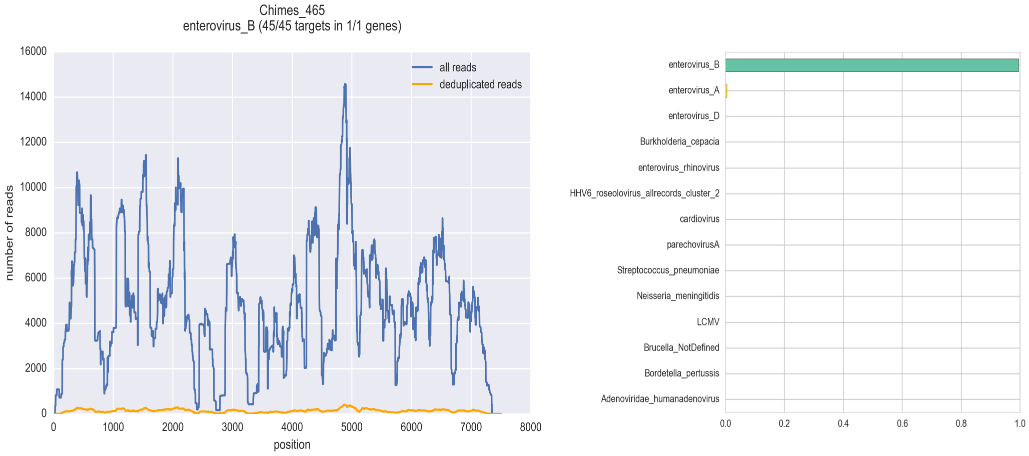


## (b)

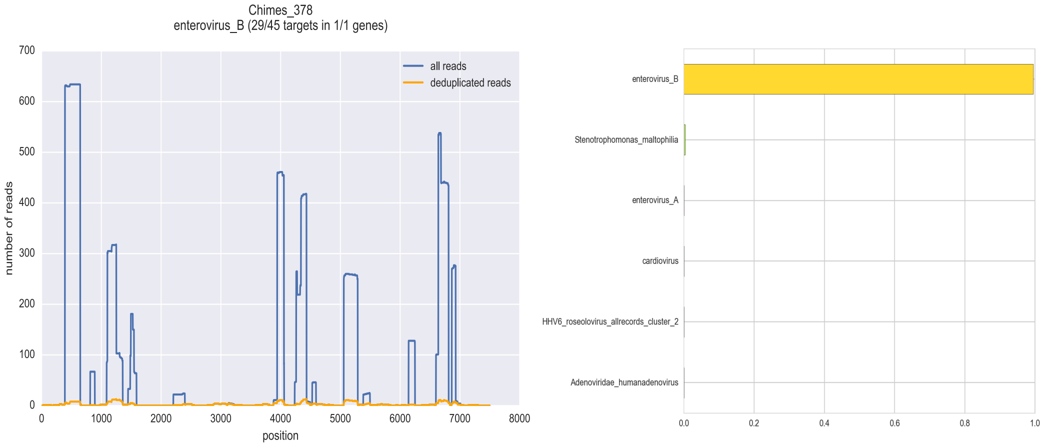

**(c)**

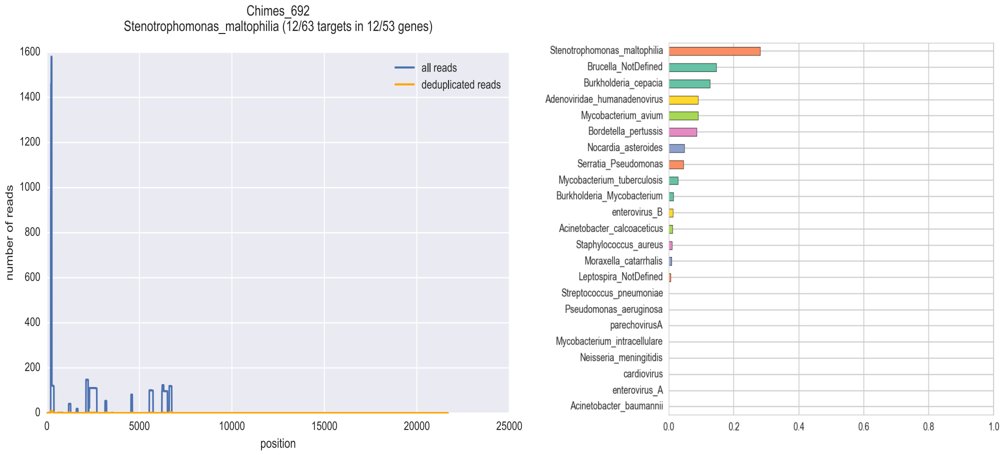


**(d)**

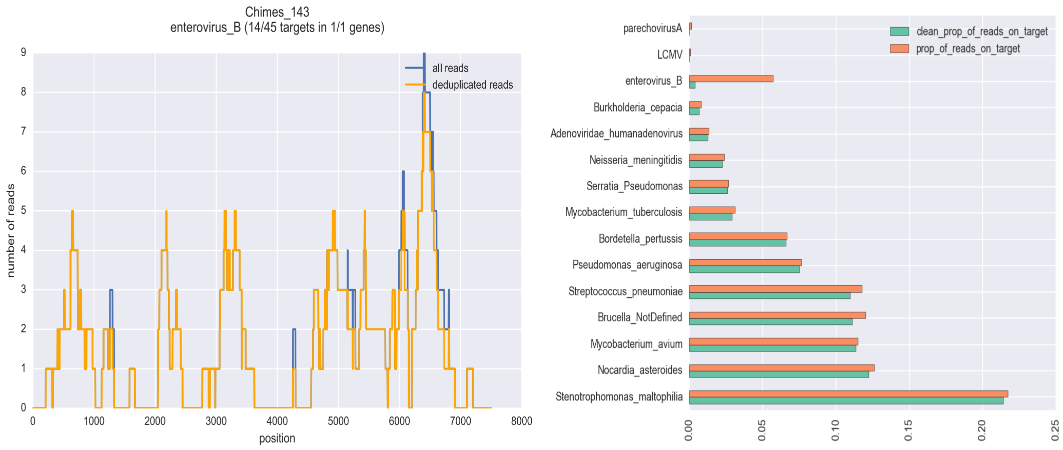


**Supplementary Figure 5.** Illustrative examples of Castanet analysis for detection of organisms in clinical samples. Left panels: Orange, deduplicated reads; blue, all reads. Right panels: proportion of all mapped reads in the sample that were mapped to this organism. (a) Strongly positive sample; (b) Weakly positive sample, showing characteristic peaks of duplicated reads resulting from capture of partial genome; (c) Negative sample; (d) Negative sample showing characteristic presence of low-level background reads due to index mis-assignment, which can then be removed bioinformatically (right panel, “clean prop of reads on target”).
